# A sensorimotor enhanced neuromusculoskeletal model for simulating postural control of upright standing

**DOI:** 10.1101/2024.03.13.584822

**Authors:** Julian Shanbhag, Sophie Fleischmann, Iris Wechsler, Heiko Gassner, Jürgen Winkler, Bjoern M. Eskofier, Anne D. Koelewijn, Sandro Wartzack, Jörg Miehling

**Affiliations:** Engineering Design, Department of Mechanical Engineering, Friedrich-Alexander-Universität Erlangen-Nürnberg, Germany; Machine Learning and Data Analytics Lab, Department Artificial Intelligence in Biomedical Engineering (AIBE), Friedrich-Alexander-Universität Erlangen-Nürnberg, Germany; Department of Molecular Neurology, Universitätsklinikum Erlangen, Friedrich-Alexander-Universität Erlangen-Nürnberg, Germany; Chair for Autonomous Systems and Mechatronics (ASM), Department of Electrical Engineering, Friedrich-Alexander-Universität Erlangen-Nürnberg, Germany

## Abstract

The human’s upright standing is a complex control process that is not yet fully understood. Postural control models can provide insights into the body’s internal control processes of balance behaviour. Using physiologically plausible models can also help explaining pathophysiological motion behaviour. In this paper, we introduce a neuromusculoskeletal postural control model using sensor feedback consisting of somatosensory, vestibular and visual information. The sagittal plane model was restricted to effectively six degrees of freedom and consisted of nine muscles per leg. Physiological plausible neural delays were considered for balance control. We applied forward dynamic simulations and a single shooting approach to generate healthy reactive balance behaviour during quiet and perturbed upright standing. Control parameters were optimized to minimize muscle effort. We showed that our model is capable of fulfilling the applied tasks successfully. We observed joint angles and ranges of motion in physiological plausible ranges and comparable to experimental data. This model represents the starting point for subsequent simulation of pathophysiological postural control behaviour.

## 1 Introduction

Human upright standing is inherently unstable. Nevertheless, the human’s postural control system is able to produce muscle forces and thereby joint torques to maintain the body in an upright position, even against external perturbations. Up to a certain degree this is possible without the need to take a correction step. One typical symptom of neurological disorders like Parkinson’s Disease (PD) is an impaired function of the postural control system which results in difficulties maintaining balance during daily tasks. A detailed understanding of the body’s internal control processes during postural control is essential to explain pathophysiological postural control and to be able to give tailored therapy recommendations to patients suffering from neurological disorders like PD.

To maintain balance, the human body continuously initiates muscle forces to keep the center of mass (COM) within the base of support (Winter, 1995). The base of support is defined by the area beneath the contact points of the feet with the ground. The central nervous system regulates information from the somatosensory, vestibular and visual systems to gain current body states and initiates suitable muscle excitations that lead to adequate muscle forces to keep the body in balance (Forbes et al., 2018). The somatosensory system consists of proprioception and cutaneous receptors. Proprioceptive information is perceived by muscle spindles and Golgi tendon organs. Muscle spindles are located in the skeletal muscles and sense muscle lengths and lengthening velocities (Kröger & Watkins, 2021). Golgi tendon organs are located at the interface between muscle and tendon and sense muscle tendon forces. Cutaneous receptors deliver tactile information about the pressure distribution underneath the feet (Jahn & Wühr, 2020), which includes changes in the location of the center of pressure (COP). The vestibular system is sensitive to linear and angular motion and orientation of the head. It consists of two structures located within the inner ear, the otolith organs and semicircular canals. Otolith organs detect linear accelerations as well as the head tilt with respect to the gravitational field, semicircular canals the rotational head accelerations (Jahn & Wühr, 2020; Mahboobin et al., 2002). The visual system provides information about the direction and speed of body sway (Jahn & Wühr, 2020). All these sensory information are centrally integrated to ensure a reliable and robust interpretation of the body state that can be used for postural control reactions. Lower level controls, like reflexes, are generated in the spinal cord, higher-level controls in the supra-spinal cord (Jahn & Wühr, 2020). This process is subject to neural delays consisting of processing, transmission and activation dynamics delays.

Biomechanical model simulations can be used to simulate and analyze postural control behaviour. Upright standing is often investigated using simplified models such as single inverted pendulum models (Goodworth & Peterka, 2009; Masani et al., 2006; Welch & Ting, 2008). More detailed models can consist of a higher number of degrees of freedom (DOF) and muscles (Kaminishi et al., 2019; Koelewijn & Ijspeert, 2020; Versteeg et al., 2016). Simulations of musculoskeletal human models often focus on proprioceptive information (Koelewijn & Ijspeert, 2020; Suzuki & Geyer, 2018) or assume the body’s full-state information to be known by the central nervous system (Welch & Ting, 2008; Yin et al., 2020). Also, the considered amount of neural delays varies a lot between the different models. An overview about different simulation approaches of postural control, which biomechanical human models and control strategies are used, is given by Shanbhag et al., 2023. To gain a detailed understanding about internal processes during postural control, it is necessary to consider all of the different sensory systems that the human body uses to maintain balance. A clear distinction between the origins of different sensory signals a model is using, like from Jiang et al., 2017, is rarely done in postural control simulations. However, detailed postural control models considering such distinctions and all sensory systems used for postural control, and also neural delays in physiologically plausible ranges, could be capable of covering many aspects of postural control and giving insights into internal processes of the body.

In this paper, we use a forward dynamic approach to simulate balance control. We introduce a postural control model for upright standing using a musculoskeletal human model with nine DOF and 18 muscles. The model considers somatosensory, vestibular as well as visual information for generating muscle feedback. Also, physiologically plausible neural delays are added within the neural circuitry, depending on muscle position and information type. The model is able to simulate quiet and perturbed upright standing.

## 2 Material and Methods

We used a generic musculoskeletal human model to conduct forward-dynamic simulations of postural control behaviour of perturbed and quiet upright standing. The simulations in this study were applied using the software framework SCONE 2.3.0 (Geijtenbeek, 2019) with Hyfydy (Geijtenbeek, 2021). The implementation consists of three elements: A musculoskeletal human model, a neural controller and an optimization of free control parameters. In the following, we present in detail the musculoskeletal model, the control design, as well as the optimization steps. Additionally, the simulation approach and the experimental data with which the simulation results are compared are described.

### 2.1 Musculoskeletal Model

For our simulations, we used a musculoskeletal human model based on Delp et al., 1990 with updates from (Rajagopal et al., 2016), distributed as part of SCONE. This version is a planar model (sagittal plane) with seven segments, consisting of trunk-pelvis, and each upper leg, lower leg and foot. The model is restricted to nine DOF, three DOF per leg (ankle, knee and hip joint) and three DOF between the pelvis and the ground. We assumed a symmetric motion behaviour. As a result, the model effectively has six DOF. Nine Hill-type muscles (Millard et al., 2013) are considered for each leg: Gluteus maximus (GLU), iliopsoas (IL), hamstrings (HAM), biceps femoris short head (BFSH), rectus femoris (RECT), vastus intermedius (VAS), gastrocnemius medialis (GAS), soleus (SOL), and tibialis anterior (TA). Values for muscle parameters were set according to Delp et al., 1990. Ground contact is modelled via two viscoelastic Hunt-Crossley contact spheres per foot, one on the heel and one on the forefoot.

### 2.2 Neural Control Design

We implemented a neural control circuit to simulate postural control behaviour of upright standing using the aforementioned musculoskeletal human model. Our aim was to establish processes on a physiological plausible basis: All sensory information that are used in the physiological process of postural control should be available for the model’s control as well. State information of the body, gained by the different sensory systems of the body, were considered to generate feedback signals, depending on their corresponding gain factors. Additionally, the feedback loop is subject to neural delays *τ* (section 2.2.2). The whole postural control model is shown in Fig. 1.

**Figure 1.**
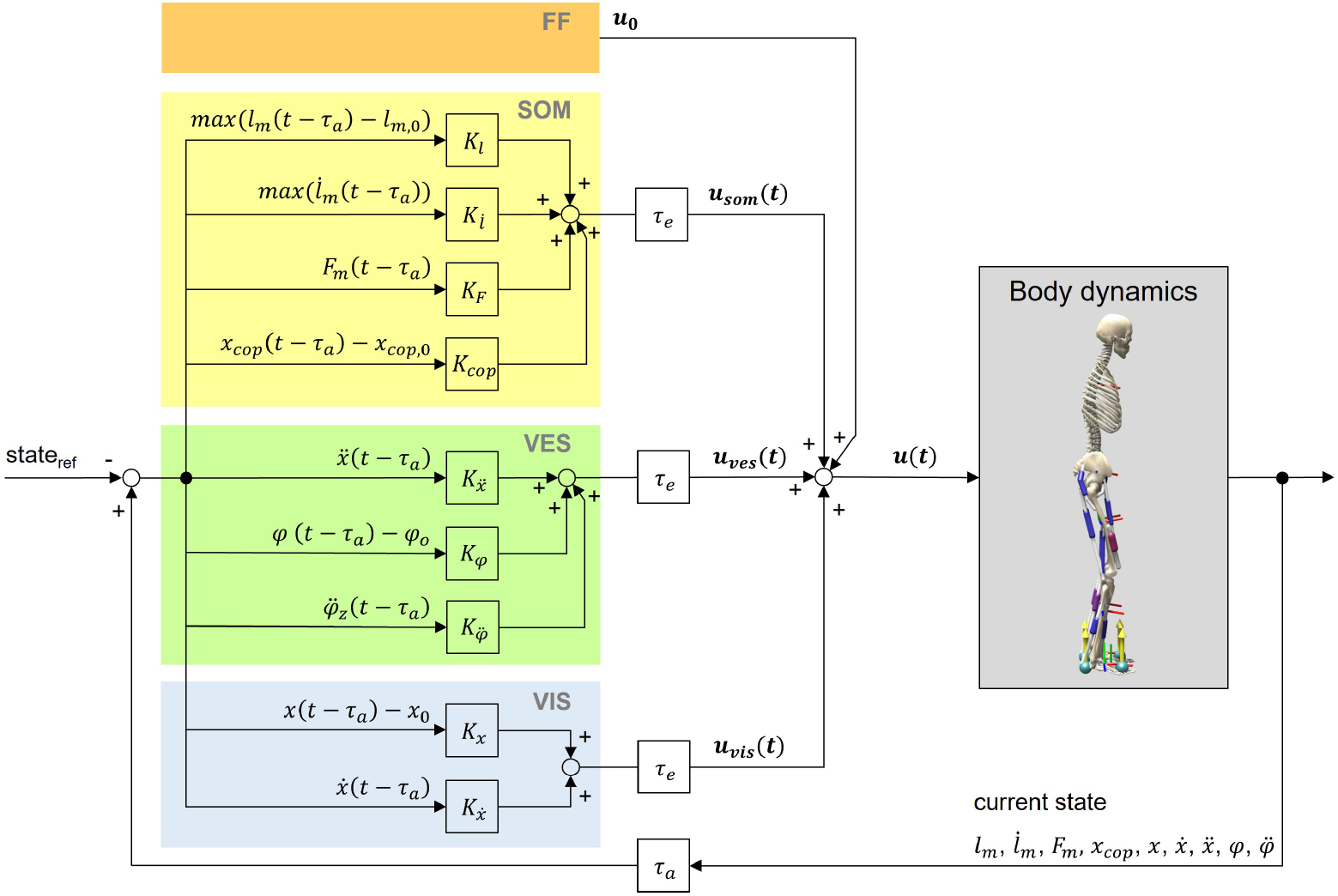
Postural control model. The neural controller generates control signals based on feedforward input (FF) as well as somatosensory (SOM), vestibular (VES) and visual (VIS) feedback. Feedback values are generated by comparing time-delayed states of the model with their corresponding reference states. Indicated delays belong to the body’s afferences (*τ*_*a*_) and efferences (*τ*_*e*_). For calculations, we used a lumped delay *τ* consisting of both signal transmission and processing delays (section 2.2.2). The reference position is an upright standing pose. Signals from the neural controller lead to muscle excitations *u*(*t*) of the musculoskeletal human model.

#### 2.2.1 Feedback controller

The total muscle excitation that is continuously calculated for each individual muscle is the sum of the different sensory systems’ feedback and is represented in equation 1:

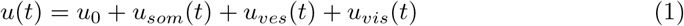

The total muscle excitation *u*(*t*) consists of a feedforward element *u*_0_, somatosensory feedback *u*_*som*_(*t*), vestibular feedback *u*_*ves*_(*t*) and visual feedback *u*_*vis*_(*t*). Equations 2-4 are used to determine the corresponding feedback elements.

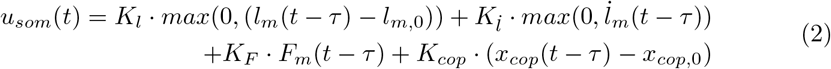

*K*_*l*_, *K*_*l*_?, *K*_*F*_ and *K*_*cop*_ are gain factors based on proprioceptive and tactile information of the somatosensory system. *l*_*m*_(*t − τ* ), *l*?_*m*_(*t − τ* ) and *F*_*m*_(*t − τ* ) represent the time-delayed normalized length, lengthening velocity and force of the corresponding muscle. *l*_*m*,0_ is the offset length of the muscle, *x*_*cop*,0_ the initial COP value and the midpoint between the two contact points of the feet with the ground.

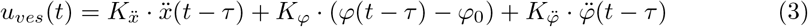

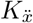, *K*_*φ*_ and 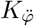 are gain factors based on vestibular information. 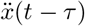 represents the time-delayed linear acceleration, *φ*(*t − τ*) the time-delayed orientation and 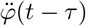 the time-delayed angular acceleration of the head with respect to the environment. *φ*_0_ is the initial orientation of the head.

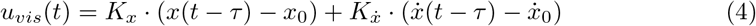

*K*_*x*_ and 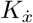 are gain factors based on visual information. *x*(*t − τ* ) and 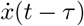 represent the time-delayed position and velocity of the head with respect to the environment.

All feedback gains, the offset muscle lengths *l*_*m*,0_ as well as feedforward excitations *u*_0_ were optimized in the optimization step (section 2.3).

Additionally, we applied a small amount of random Gaussian noise to the model’s sensor systems as well as to the actuators assuming that these systems do not work noise free in the human body, consisting of base noise and noise depending on the signal amplitude *s*, which is either sensor information or muscle excitation.

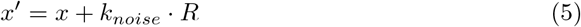

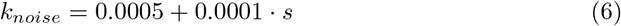

x is the true signal, *k*_*noise*_ the signal-dependent noise amplitude and *R* a randomly generated Gaussian distributed number. *x*^*′*^ represents the resulting signal’s value.

#### 2.2.2 Neural delays

Depending on the muscle position in the body and the sensory information type, we considered different amounts of neural delays. For each muscle we defined a lumped neural delay *τ* consisting of transmission and processing delay. We set neural delays according to assumptions of Li et al., 2012. For muscle reflexes based on proprioceptive information reactions up to 25 ms are possible. Depending on the muscle’s position, the reflex delay can increase by additional 25 ms due to the longer transmission distance from the central nervous system to shank muscles compared to hip muscles. For vestibular and visual information a higher amount of processing is necessary. Therefore, additional 100 ms were assumed for each muscle. All neural delays that we used are summarized in Tab. 1.

**Table 1.** Neural delays for each muscle depending on the muscle’s position and the sensory information type.

| Muscle | Neural delays |  |  |
| --- | --- | --- | --- |
|  | Somatosensory<br>information | Vestibular<br>information | Visual<br>information |
| Gluteus maximus | 25 ms | 125 ms | 125 ms |
| Iliopsoas | 25 ms | 125 ms | 125 ms |
| Rectus femoris | 25 ms | 125 ms | 125 ms |
| Hamstrings | 25 ms | 125 ms | 125 ms |
| Biceps femoris short head | 35 ms | 135 ms | 135 ms |
| Vastus intermedius | 35 ms | 135 ms | 135 ms |
| Gastrocnemius medialis | 50 ms | 150 ms | 150 ms |
| Soleus | 50 ms | 150 ms | 150 ms |
| Tibialis anterior | 50 ms | 150 ms | 150 ms |

### 2.3 Optimization of Control Parameters

The control gains, offset muscle lengths and feedforward excitations were optimized using single shooting and the pre-implemented covariance matrix adaption evolution strategy (CMA-ES) algorithm (Igel et al., 2007) in SCONE. An optimization was solved to find control parameters to minimize muscular effort and to fulfill additional constraints, such as knee and hip joint limits and keeping the model’s COM higher than 60% of the initial height. Effort minimization is chosen as this is assumed to be the objective of the central nervous system when creating movements (Selinger et al., 2015). This leads to the following optimization problem:

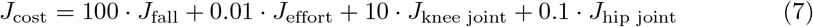

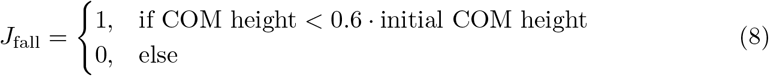

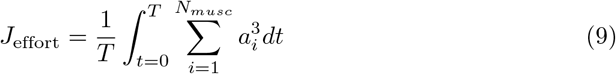

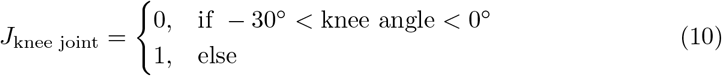

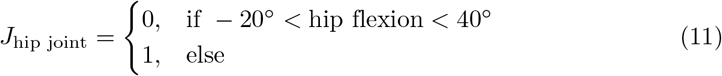

*N*_*musc*_ represents the number of muscles and *a*_*i*_ the activation of each muscle. For each scenario, six optimizations with different random seeds were performed. Optimizations ended as soon as the averaged reduction of the cost function’s result was smaller than 1e-5 compared to the previous iteration.

In this model, we assumed a symmetric postural control behaviour, so left and right muscle excitations are calculated identically. For parameters’ initial guesses, we considered findings of Peterka, 2018 where sensory feedback weightings were identified. They reported postural control to be based 50% on proprioceptive, 33% on visual, and 17% on vestibular information for low amplitude perturbations. We applied this ratio to our feedback gains’ initial guesses, where applicable. Boundaries for parameter optimization were defined as -3 and 3 for control gains, as 0.001 and 0.2 for feedforward excitation *u*_0_, and as 0.1 and 2 for the muscle’s offset length *l*_*m*,0_.

### 2.4 Simulation Approach

Full simulations lasted 75 seconds. Because the first seconds are not representative due to a manually predefined starting position, we evaluated simulation results from 15 seconds simulation time for 60 seconds in total. Simulation frequency was set to 200 Hz.

The model’s starting position was always an upright stance (pelvis tilt: -10.0°, hip flexion: 20.0°, knee angle: -20°, ankle angle: 10°) and adopted from the pre-defined SCONE settings (Geijtenbeek, 2019). To evaluate our postural control model, we tested it under different scenarios: In the first step, we simulated an quiet standing (no external perturbations). Results were compared to experimental data. Additionally, we tested an upright standing on a moving platform. We applied translational perturbations to the platform comparable to the study of Wang and van den Bogert, 2020. This resulted in anterior posterior perturbations of the model and allowed us to compare our simulation results with the experimental results of this open accessible data set.

### 2.5 Experimental Data

We conducted a study to measure postural control behaviour during quiet upright standing. 8 healthy participants (4 male, 4 female, age: 51.63 ± 23.12 years) were included in this study. All participants performed an upright standing task with eyes open for 60 seconds. They were instructed to place their feet shoulder-width apart and focus on a sign at eye level in front of them. Hands were placed on the hips during this task. We collected the data in a motion laboratory using an optical motion capture system (VICON Vero, ten cameras, 100 Hz) and two force plates (AMTI, 1000 Hz). We recorded the data with 45 reflective body markers according to the Plug-in Gait model (Vicon Motion Systems Limited UK, 2021), with four additional markers on each medial knee and ankle. These four medial markers were detached after an initial calibration measurement. For processing the experimental data, we used OpenSim 4.4 (Seth et al., 2018) and a generic three-dimensional model, based on Delp et al., 1990. The model consisted of 17 DOF and 8 segments. The processing itself was conducted using the tool AddBiomechanics (Werling et al., 2023) that includes an automatic model scaling and inverse kinematics from human motion data. In the end, results were filtered with a third order Butterworth filter with a cutoff frequency of 6 Hz.

Experimental data that we used to compare an upright standing on a moving platform (Wang & van den Bogert, 2020) was collected from 6 participants. Each of them fulfilled two perturbed standing tasks. Measurement data was processed using a two-dimensional sagittal plane model (three DOF) consisting of hip, knee and ankle joint. During calculations, left and right joint angles were averaged, as movements were assumed to be symmetrical.

### 2.6 Data evaluation

To evaluate our postural control model for upright standing, we compared simulation results with experimental data (section 2.5). We used own collected experimental data to evaluate the unperturbed scenario. For the upright standing on a moving platform, we compared simulation results with the publicly available data set of Wang and van den Bogert, 2020. In the dataset records, platform movement started after approximately 12 seconds. For evaluation, we used the time period between 15 seconds and 75 seconds for both our simulation and experimental data to compare areas with a high amount of perturbation. For all scenarios we analyzed joint angles and COP values over 60 seconds.

We averaged experimental data to compare them with simulation results. We calculated root mean square errors (RMSEs) of simulations and experimental data as well as ranges of motion (ROMs) of simulations and average ROMs for experimental data. To calculate RMSEs, we used experimental data of each participant averaged for each time step resulting in average time courses. To gain average ROMs, we averaged all participant’s individual ROMs. Because the absolute COP position depends on the definitions of the force plates’ coordinate systems and standing positions of participants, we compared just the COPs range.

## 3 Results

The simulation model was able to successfully fulfill the unperturbed standing task for the complete simulation time. Fig. 2 shows joint angles of the simulation and each participant of the experiments. We observed RMSEs of 5.28^*°*^ (pelvis tilt), 4.93^*°*^ (hip angle), 1.54^*°*^ (knee angle) and 0.91^*°*^ (ankle angle) between simulations and average experimental data. Simulation results show joint angles’ ROMs of 0.63^*°*^ (pelvis tilt), 0.72^*°*^ (hip angle), 1.16^*°*^ (knee angle) and 0.67^*°*^ (ankle angle). Average experimental data show ROMs of 2.59 ± 0.66^*°*^ (pelvis tilt), 2.52 ± 1.16^*°*^ (hip angle), 1.38 ± 0.54^*°*^ (knee angle) and 1.11 ± 0.30^*°*^ (ankle angle). The COP range was 11.96 mm during the simulation, 25.62 ± 8.65 mm during experiments. Tab. 2 summarizes RMSEs and ROMs of joint angles and the COP of each simulation and average experimental data.

**Table 2.** Joint angles’ and COP’s parameters of the simulation and average experimental data during quiet upright standing.

| Parameters | ROM simulation | ROM experiments | RMSE |
| --- | --- | --- | --- |
| Pelvis angle [deg] | 0.63 | $2.59 \pm 0.66$ | 5.28 |
| Hip angle [deg] | 0.72 | $2.52 \pm 1.16$ | 4.93 |
| Knee angle [deg] | 1.16 | $1.38 \pm 0.54$ | 1.54 |
| Ankle angle [deg] | 0.67 | $1.11 \pm 0.30$ | 0.91 |
| COP [mm] | 11.96 | $25.62 \pm 8.65$ | N/A |

**Figure 2.**
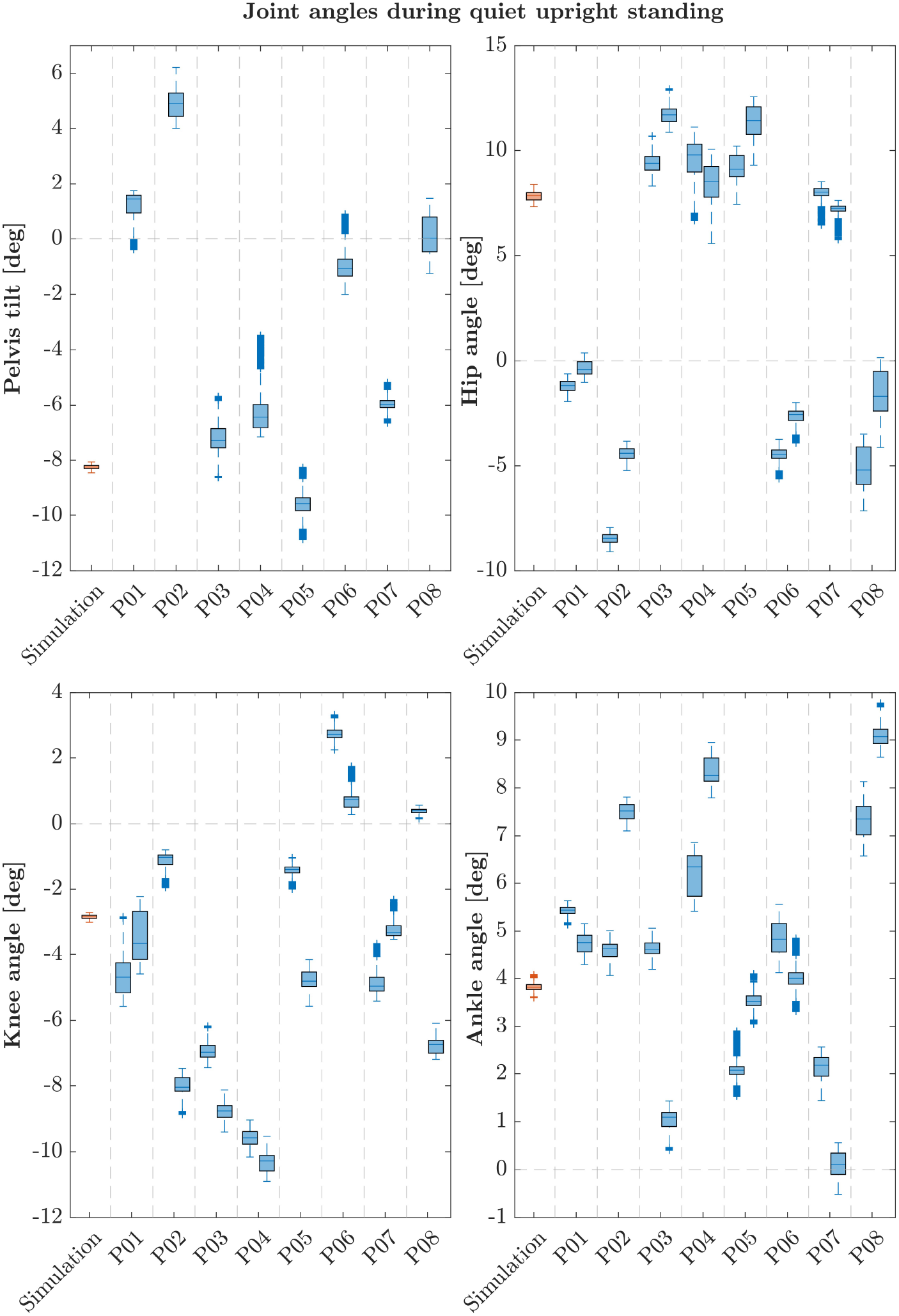
Box-plots showing median and interquartile ranges of joint angles during quiet upright standing of simulation and each participant. Simulations assumed left and right symmetry, experimental data show left and right joint angles of each participant separately.

The postural control model was also capable of maintaining balance on a moving platform. We compared simulation results with the published joint angles (Wang & van den Bogert, 2020). Fig. 3 shows joint angles of simulation and each participant of the experiments. As there was no pelvis tilt given in the published dataset, we only compared hip, knee and ankle angles. Resulting joint angle courses of simulation and average experimental data are represented in Fig. 4, COP courses in Fig. 5. We observed RMSEs of 11.11^*°*^ (hip angle), 1.86^*°*^ (knee angle) and 1.04^*°*^ (ankle angle) between simulation and experiments. Simulation results show joint angles’ ROMs of 8.84^*°*^ (pelvis tilt), 8.29^*°*^ (hip angle), 8.73^*°*^ (knee angle) and 4.16^*°*^ (ankle angle). Average experimental data show ROMs of 23.46 ± 14.97^*°*^ (hip angle), 13.34 ± 8.35^*°*^ (knee angle) and 9.10 ± 5.47^*°*^ (ankle angle). The COP range was 124.77 mm during the simulation, 120.43 ± 25.55 mm during experiments. Joint angles’ RMSEs, ROMs and ranges of the COP are summarized in Tab. 3.

**Table 3.** Joint angles and COP’s parameters of the simulation and average experimental data during upright standing on a moving platform. No pelvis angles were given for experimental results.

| Parameters | ROM simulation | ROM experiments | RMSE |
| --- | --- | --- | --- |
| Pelvis tilt [deg] | 8.84 | N/A | N/A |
| Hip angle [deg] | 8.29 | $23.46 \pm 14.97$ | 11.11 |
| Knee angle [deg] | 8.73 | $13.34 \pm 8.35$ | 1.86 |
| Ankle angle [deg] | 4.16 | $9.10 \pm 5.47$ | 1.04 |
| COP [mm] | 124.77 | $120.43 \pm 25.55$ | N/A |

**Figure 3.**
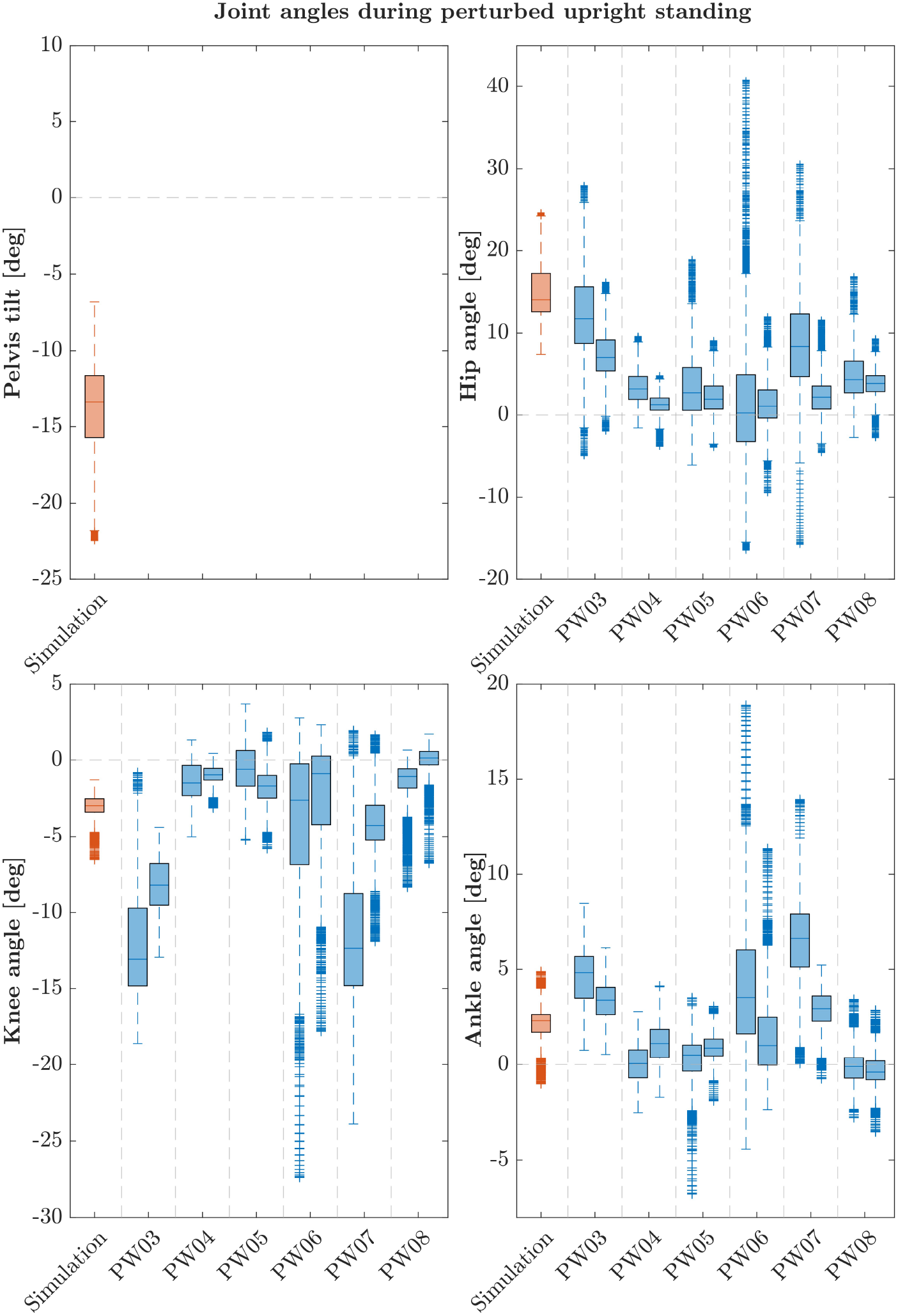
Box-plots showing median and interquartile ranges of joint angles during upright standing on a moving platform of simulation and each participant. During the experiments, each subject fulfilled the perturbed standing task twice. Note that these participants are not the same as in the quiet standing scenario.

**Figure 4.**
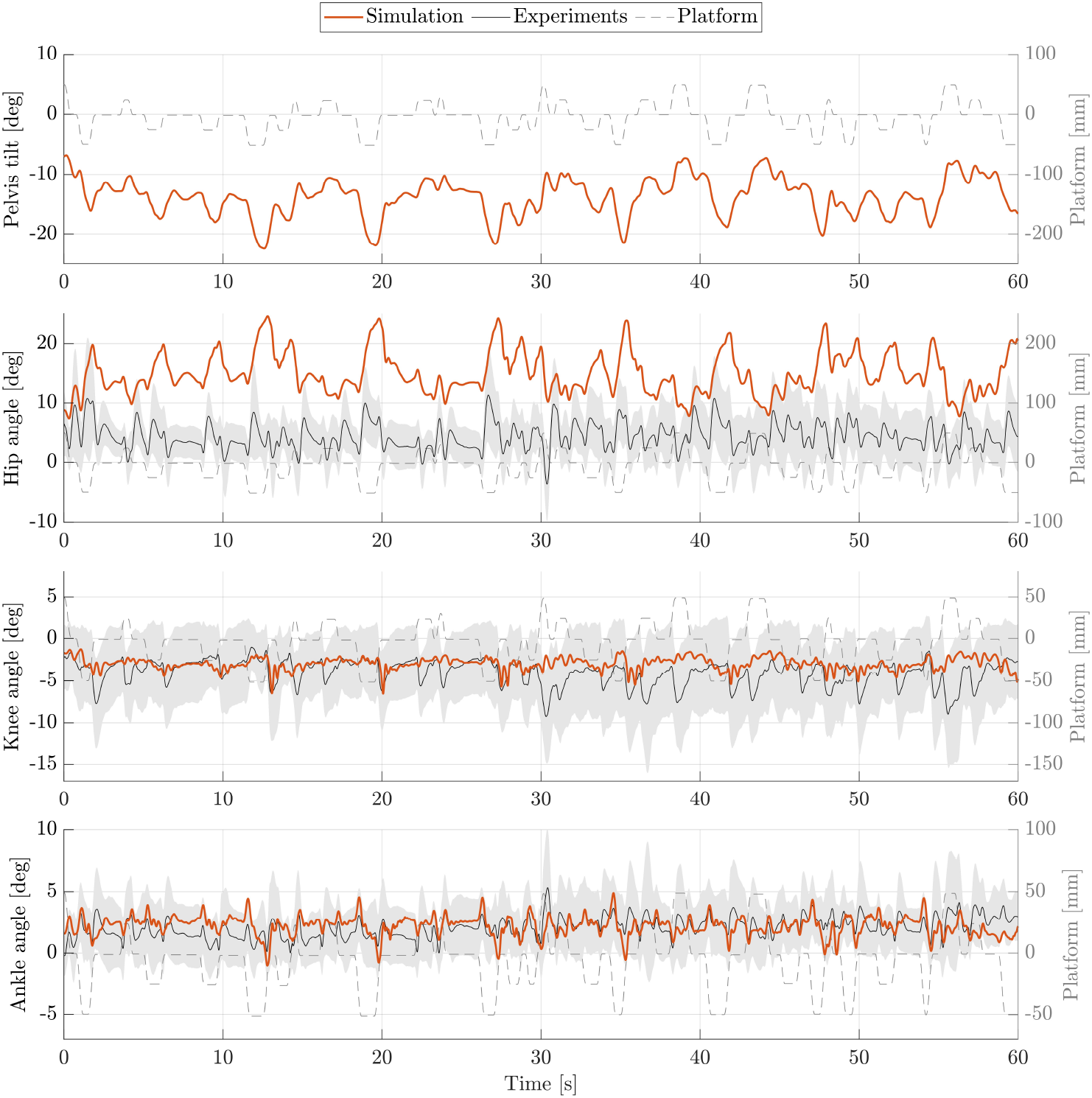
Joint angles during upright standing on a moving platform. Joint angles of simulation (orange) and experiments (black) are shown for 60 seconds.

**Figure 5.**
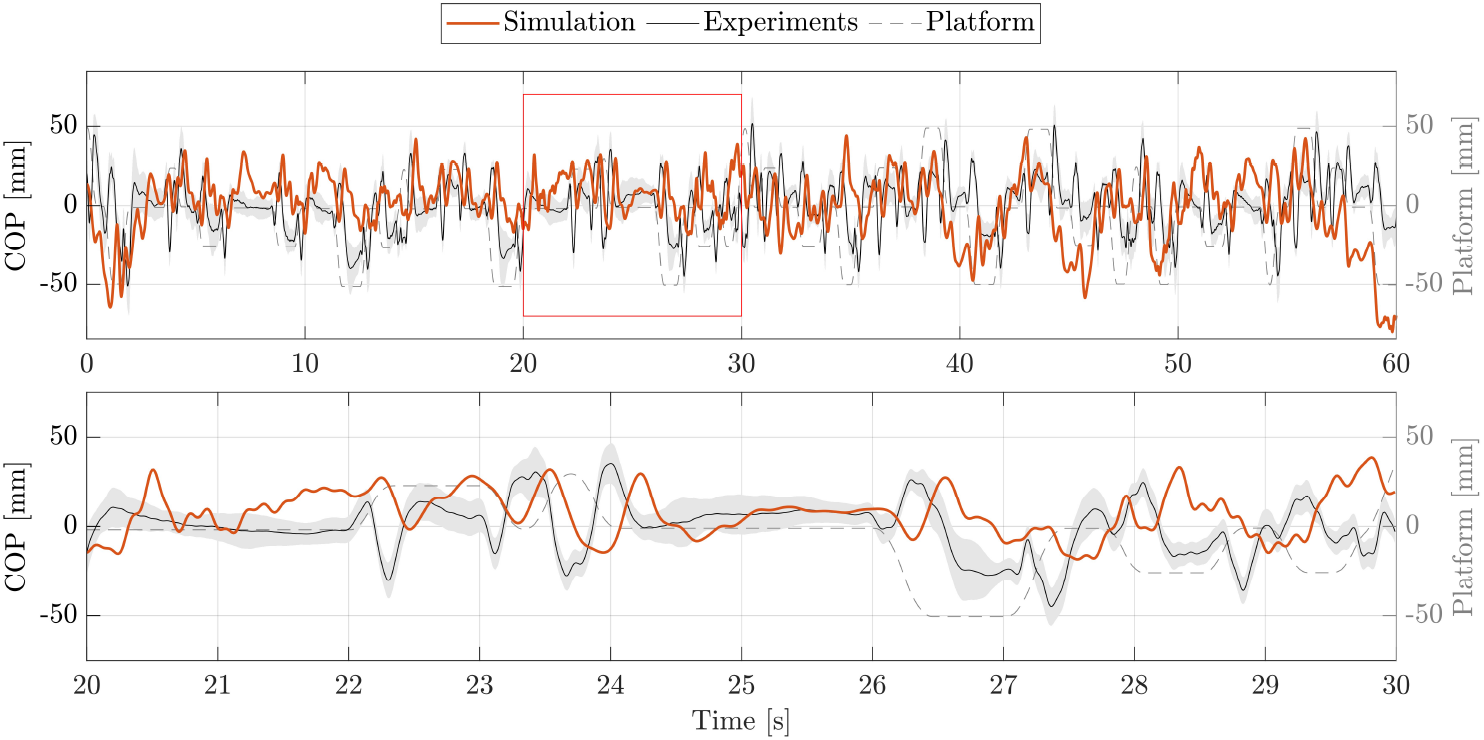
COP during upright standing on a moving platform. The COP of simulation (orange) and experiments (black) is shown for 60 seconds. Additionally, an exemplary section of 10 seconds (highlighted with a red frame) is represented as zoom-in below.

## 4 Discussion

In this paper, we aimed to simulate postural control behaviour using a musculoskeletal human model and complex sensor feedback of all sensory systems involved in postural control considering physiologically plausible neural delays. Compared to many other postural control models, our model uses somatosensory, vestibular as well as visual information for balance control under the influence of neural delays in physiologically plausible ranges. The model is able to fulfill an upright standing task in unperturbed and perturbed situations. Motion behaviour shows to be comparable to experimental data of healthy participants for both simulation scenarios. For both, the unperturbed as well as the perturbed situation, the model’s motion is comparable to the ones observed during experimental measurements.

We observed RMSEs smaller than 1.9^*°*^ for ankle and knee angles, only hip and pelvis showed a higher variation between simulation and experiments (4.93-11.11^*°*^). This is because the stable standing pose that was found by the optimization, especially during the standing on a moving platform, consists of a higher hip flexion and a more forward leaning torso. As the initial standing pose has influences on the reference control parameters, a different initial pose could improve the torso orientation. Also, the model’s reference COP position is currently defined as the midpoint between the two contact spheres of each foot. In reality, the COP might not lie directly in this point. As we could not compare absolute COP positions, simulations and experiments could differ in this aspect. During quiet standing, ROMs are comparable, only the COP range shows to be considerably smaller during the simulation. During the perturbed standing, the simulation’s ROM is noticeable smaller for all joint angles, the COP range is comparable to the experiments. It has to be said, that also the experimental data itself showed high standard deviations of the ROMs. This means, that the amount of ROM varies widely between the individual participants. Also, experimental results showed high variation in absolute mean values in general which is shown in Fig. 2 and 3. Considering this, simulation results show realistic motion behaviour.

Besides internal noise of sensory systems and muscle actuators, also low-frequent disturbances caused by breathing or the heart beat influence motion during postural control (Forbes et al., 2018). These effects are currently not considered in our model and could explain the remaining difference between simulation and experimental ROMs.

We used a sagittal plane musculoskeletal human model to simulate postural control. We are aware that not only anterior-posterior, but also medio-lateral movements are relevant in order to holistically simulate motion behavior. Additionally, our current simulation approach creates symmetrical motion. In reality, human motion is never completely symmetrical. Still, a sagittal plane model will already be capable of providing some insights into impaired neural control as well, even if it is not complete. We assumed different neural delays depending on muscle position and sensor information type. It has to be mentioned that the specific amount of different neural delays is still being discussed. In this respect, current models for postural control differ considerably, like with 100 ms (van der Kooij et al., 2001), 120 ms (Jiang et al., 2017), 150 ms (Van Wouwe et al., 2022) or 185 ms (Masani et al., 2006). However, at the same time, postural control models could help to identify neural delays by investigating differences in motion behaviour resulting from adaptions of neural delays. Our model assumes a multisensory integration in form of a weighted sum of several sensory information in the feedback loop. Also this internal process of sensory integration of the body is still unclear. Many models use this approach to fuse sensor information (Goodworth & Peterka, 2009; Jiang et al., 2017; Van Wouwe et al., 2022). Other models use for example optimal estimator methods (Kuo, 2005; van der Kooij et al., 2001) to process the information before initiating corresponding model reactions. Some studies inform about frequency-dependencies of the body’s sensory systems (Forbes et al., 2018; Jahn & Wühr, 2020; Peterka, 2018). Optimal estimator methods can take this aspect into account. In our approach, this characteristic is currently not addressed. We aimed to simulate the reactive postural control behaviour of upright standing. Up to this point, our model considers aspects of spinal and supra-spinal control. Motion aspects such as voluntary movements, are currently not included in the model.

Postural control behaviour can differ substantially between subjects. Even in healthy individuals, factors like age influence postural control significantly (Rinaldi et al., 2009; Van Humbeeck et al., 2023). Even though we ensured to include subjects of different sex and age, a higher number of subjects could still influence our reference values. We used experimental data of 8 subjects (age: 51.63 ± 23.12 years) to compare them with our simulation results and gain an overall impression of our model compared to human data. In a next step, also ageing effects could be considered to model age-specific postural control. Up to this point, our simulations were conducted with a generic OpenSim model. A next step could be to use personalized musculoskeletal human models to get a more subject-specific motion behaviour.

Additionally, it is important to keep in mind that even though if a postural control model shows human-like motion behaviour, this does not necessarily prove that the model mimics the control processes of real humans. Nevertheless, these models may help to gather insights on the differences between physiological and pathophysiological control.

## 5 Conclusion and Outlook

In this paper, we introduced a musculoskeletal postural control model using complex sensor feedback consisting of somatosensory, vestibular and visual information considering physiologically plausible neural delays. It is able to maintain balance in both unperturbed as well as perturbed scenarios. The simulated motion behaviour showed to be comparable to empirical data of healthy participants.

This model will serve as a basis to simulate and even characterize motion behaviour of persons suffering from neurological disorders like PD. In a next step, parameters that are thought to be the cause of symptoms such as postural control impairments could be adjusted. This will allow us to further investigate the balance behavior of Parkinson’s patients and to assess, for example, the effects of different rehabilitation interventions.

## Acknowledgments

The Ethics Committee of the FAU Erlangen-Nürnberg approved the protocol of this prospective study and the research was conducted in accordance with the Declaration of Helsinki. All participants have signed a written consent form for this study.

